# Impaired AMPK activity contributes to the inflammatory phenotype and the reduced phagocytosis capacity of VASP-deficient macrophages

**DOI:** 10.1101/2023.04.19.537577

**Authors:** Hebatullah Laban, Timo Frömel, Ingrid Fleming, Peter M. Benz

## Abstract

Macrophage polarization plays an important role in tissue regeneration. Numerous factors and signaling molecules affect polarization processes. Here we investigated the consequences of the genetic deletion of vasodilator-stimulated phosphoprotein (VASP), which increases macrophage M1 polarization through augmented signal transducer and activator of transcription 1 (STAT1) signaling, and AMP-activated protein kinase (AMPK), which attenuates inflammation by inhibiting STAT1 expression and signaling. While a basal activity of AMPK (phosphorylation on Thr172) was detected in macrophages from wild-type mice, AMPK phosphorylation was significantly reduced in VASP-deficient M1 macrophages *in vitro* and the expression of the pro-inflammatory cytokines TNFα and IL-1β was increased in these cells. Consistent with the role of AMPK in macrophage phagocytosis, VASP^-/-^ macrophage phagocytosis was also significantly impaired. Interestingly, impaired phagocytosis could be rescued by exogenous activation of AMPK. Mechanistically, we found that VASP binds directly to protein phosphatase 1 regulatory subunit 6 (PP1-R6) and we hypothesize that VASP-binding to PP1- R6/PP1 limits the PP1-dependent de-phosphorylation of AMPK in wild-type cells. Conversely, AMPK dephosphorylation by the PP1-R6/PP1 complex is enhanced in the absence of VASP. In summary, we have identified a link between VASP and AMP-activated protein kinase (AMPK) activity, which may contribute to the pro-inflammatory phenotype of VASP-deficient macrophages.

## Introduction

The attraction of leukocytes to the injured and hypoxic tissue by chemokines and adhesion molecules plays an important role in tissue regeneration (Griffith et al., 2014; Shireman, 2007). Once in the extravascular space, leukocytes release a range of chemokines to initiate a positive feedback loop and recruit more leukocytes (Griffith et al., 2014). In addition to increasing cell infiltration, macrophage polarization is also crucial for post-ischemic vascular remodeling. Many types of macrophage polarization have been described over the last several years (Mosser and Edwards, 2008) and a classically activated or pro-inflammatory (M1) phenotype, that initially occurs during inflammation, can be reproduced *in vitro* using lipopolysaccharide (LPS), and interferon γ (IFN-γ). Skewing macrophage function towards the M1 phenotype via signal transducers and activators of transcription 1 (STAT1) and nuclear factor κB (NF-κB) is characterized by high expression levels of inducible nitric oxide synthase (iNOS), tumor necrosis factor-α (TNF-α), interleukin 1-β (IL-1β), vascular endothelial growth factor (VEGF), CD80 and other proteins (Liu et al., 2017; Sica and Mantovani, 2012).

Proteins of the Enabled/vasodilator-stimulated phosphoprotein (Ena/VASP) family are crucial mediators of cytoskeletal control, linking serine/threonine and tyrosine kinase signaling pathways to actin assembly. The N-terminal Ena/VASP homology 1 (EVH1) domain of the proteins binds to so-called FPPPP motifs, proline-rich peptides with the core consensus sequence FPxφP (where x denotes any amino acid and φ a hydrophobic residue (Niebuhr et al., 1997; Renfranz and Beckerle, 2002). The first FPPPP ligand identified was the surface protein ActA from the bacterial pathogen Listeria monocytogenes, which recruits Ena/VASP proteins to promote actin polymerization on the bacterial surface and thereby drive bacterial propulsion (Chakraborty et al., 1995; Cossart and Bierne, 2001; Niebuhr et al., 1997; Skoble et al., 2001). Subsequently, a number of functional ActA-like repeats have also been identified in host cell proteins that regulate the subcellular targeting of Ena/VASP proteins to sites of high actin turnover (reviewed in (Ball et al., 2002; Faix and Rottner, 2022)). However, the core motif alone is not sufficient for mature binding and core-flanking epitopes are required to achieve the necessary specificities and affinities. For example, the EVH1 binding affinity of the ActA peptide D**FPPPP**TDE<u>EL</u> was reduced over 100-fold by truncation of core-flanking residues to generate a **FPPPP**T peptide (Prehoda et al., 1999) and in particular the <u>EL</u> di-peptide at the C-terminal end of the ActA motif contributed to the binding (Ball et al., 2002).

As its name suggests, VASP is regulated by phosphorylation and was originally identified and isolated as a target of the cAMP/cGMP-dependent protein kinases (PK), PKA and PKG, in platelets (Halbrugge and Walter, 1989; Waldmann et al., 1987) and later in many other cardiovascular cells (Benz et al., 2009). VASP phosphorylation is important for its subcellular targeting and actin modulating properties (Benz et al., 2009). Since then, it has become clear that VASP can also be phosphorylated by other kinases including the energy sensing kinase; AMP-activated protein kinase (AMPK) (Blume et al., 2007) and dephosphorylated by protein phosphatase (PP)-1, PP2A, PP2B and PP2C (Abel et al., 1995), resulting in complex and dynamic phosphorylation patterns.

VASP is a key regulator of leukocyte recruitment and polarization in postischemic revascularization. Blood flow recovery after hindlimb ischemia, which involves angiogenesis and arteriogenesis, is accelerated in VASP^−/−^ mice and associated with increased leukocyte infiltration into ischemic muscles. VASP deficiency also altered the polarization of infiltrating macrophages through activation of signal transducer and activator of transcription 1 (STAT1) signaling, and increased pro-inflammatory cytokine generation. Indeed, blocking STAT1 activation blunted the increased cytokine expression in VASP^-/-^ M1 macrophages (Laban et al., 2018). How exactly VASP regulates STAT1 signaling is not known. Macrophage polarization goes hand in hand with metabolic reprogramming and an increase in oxidative phosphorylation. Given that VASP is a target of AMPK, we studied the role of AMPK in determining the phenotype of VASP-deficient macrophages.

## Results and discussion

Given the functional link between VASP and AMPK, we investigated whether altered AMPK activity might underlie the increased STAT1 signaling in VASP-deficient macrophages. AMPK is an important regulator of macrophage polarization and leukocyte recruitment during inflammatory responses (O’Neill and Hardie, 2013; Sag et al., 2008). Structurally, AMPK is a heterotrimeric complex comprising a catalytic α-subunit and a regulatory β- and γ- subunits. AMPK α1-subunit is the predominant isoform in immune-effector cells such as macrophages (Yang et al., 2010). The phosphorylation of Thr172 of the α-subunit activates the kinase (Jeon, 2016) and AMPK activation inhibits STAT1 expression and NF- κB signaling (Meares et al., 2013; Salminen et al., 2011). Thus, impaired AMPK activity may constitute the missing link to elevated STAT1 expression in VASP-deficient macrophages. Indeed, we observed that AMPKα phosphorylation non Thr172 was significantly attenuated in M1-polarized VASP^-/-^ deficient macrophages (Figure 1A and B). This fits well with the fact that M1 polarization is accompanied by a metabolic reprogramming and increased glycolysis (Shen et al., 2023). The lack of VASP also resulted in increased IL-1β and TNFα expression compared to wild-type cells *in vitro* (Figure 1C and D) and *in vivo* (Laban et al., 2018). In fact, AMPK has been shown to limit NF-κB signaling and thereby reduce IL- 1β and TNFα expression, potentially through the induction of activating transcription factor-3 (ATF3) (Kim et al., 2014; Sag et al., 2008; Xiang et al., 2019). However, AMPK has also been shown to attenuate inflammation by inhibiting STAT1 expression and signaling (He et al., 2015; Meares et al., 2013), consistent with the increased STAT1 expression levels reported in VASP-deficient macrophages (Laban et al., 2018). As AMPK activation enhances the phagocytic capacity of macrophages and neutrophils (Bae et al., 2011; Gregoire et al., 2018; He et al., 2019; Liu et al., 2022; Mounier et al., 2013), the phagocytic capacity of VASP-deficient macrophages was assessed. Wild-type and VASP^-/-^ bone marrow-derived macrophages were incubated with pH-sensitive fluorogenic bioparticles and relative fluorescence intensity was measured using live cell imaging. This revealed that VASP^-/-^ macrophages displayed significantly reduced phagocytic capacity compared to wild-type controls (Figure 1E and F). Interestingly, activation of AMPK with berberine increased phagocytosis in VASP-deficient cells to a level that was statistically indistinguishable from wild-type macrophages (Figure 1E and F). While other investigators have also reported a reduced phagocytic capacity of VASP^-/-^ macrophages, this phenomenon was attributed to a direct effect of VASP on the phagocytosis-induced reorganization of actin filaments in the phagocytic cup (Coppolino et al., 2001; Korber and Faix, 2022). The data shown here, however, suggest that impaired AMPK activity in VASP-deficient macrophages may, at least in part, explain the decreased phagocytosis. A link between VASP and AMPK phosphorylation in macrophages has not been reported to date. However, absence of VASP has previously been shown to reduce AMPK activation and hepatic fatty acid oxidation. Interestingly, restoring AMPK activity by AICAR treatment in vivo rescued the liver phenotype in VASP-deficient mice (Tateya et al., 2013).

**Figure 1.**
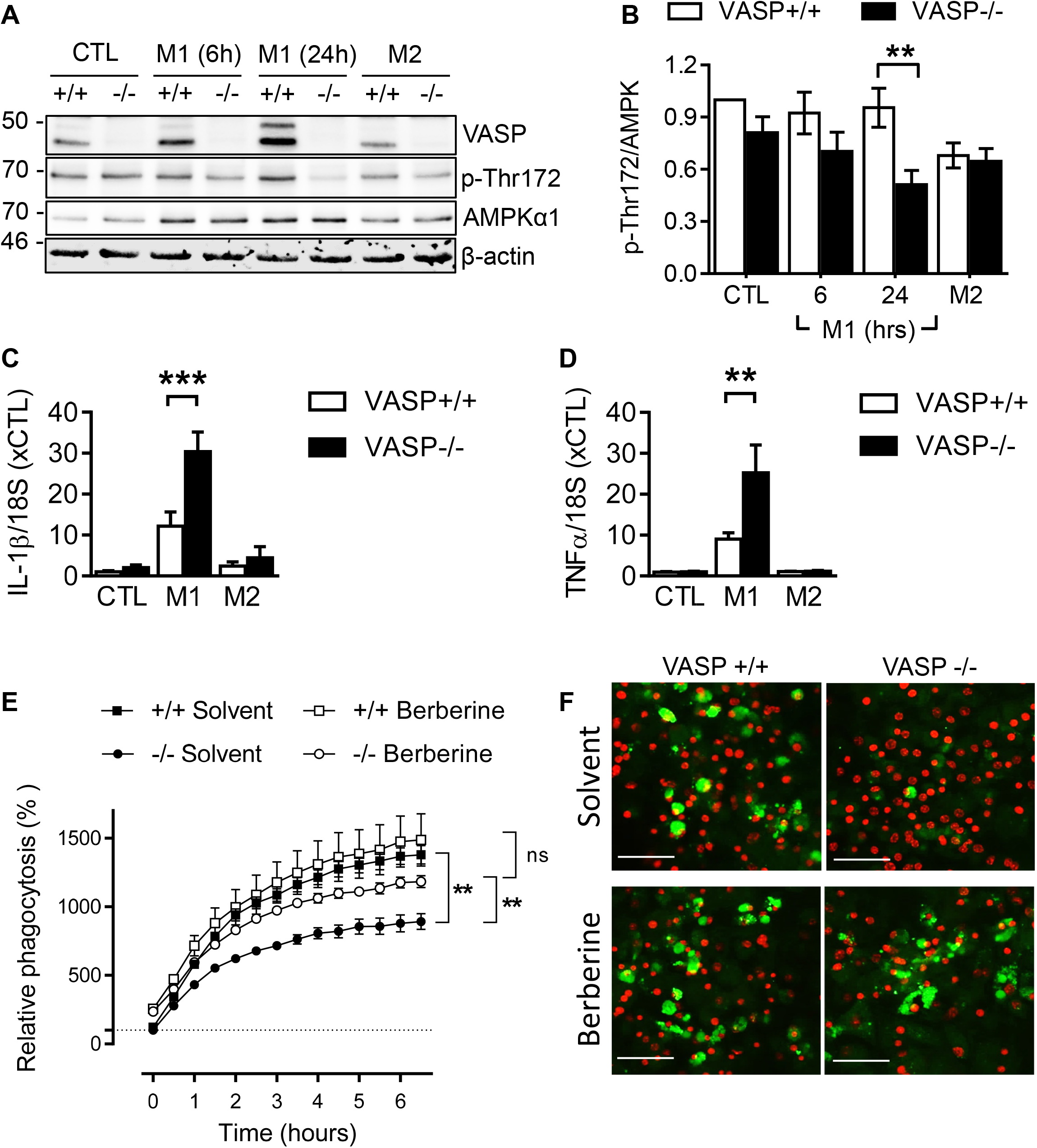
**(A, B) VASP-deficiency impairs AMPK phosphorylation**. (**A**) Representative Western blots showing the expression and Thr172-phosphorylation of AMPK in wild-type (+/+) and VASP-deficient (-/-) macrophages. Macrophages were polarized into M1 (10 ng/ml LPS + 1 ng/ml IFNγ for 6 or 24 hours) and M2 (25 ng/ml IL-4 for 24 hours). β-actin was used as loading control. (**B**) Densitometric scans were used to quantify the pThr-172 AMPK/total AMPKα1 ratio. n=7, two-way ANOVA, Bonferroni posttests; **P<0.01; wild-type *vs*. VASP-KO. **(C, D) VASP deletion increases M1 macrophage polarization in vitro**. Expression of IL-1β (**C**) and TNFα (**D**) in macrophages from WT (+/+) and VASP−/− (−/−) mice after in vitro polarization to M1 with LPS (10 ng/ml) and IFNγ (1 ng/ml) for 24 h or to M2 with IL-4 (25 ng/ml) for 24 h; n = 12 cell batches per group. Error bars, SEM; two-way ANOVA/Bonferroni; **P < 0.01; ***P < 0.001. **(E, F) Impaired VASP**^**-/-**^ **macrophage phagocytosis is rescued by AMPK activation**. (**E**) Phagocytosis assay showing the ability of BM-derived macrophages from wild-type (+/+) and VASP-deficient (-/-) mice to take up fluorogenic bioparticles after simultaneously treated with solvent or berberine (10 µmol/L, pre-incubation for 1 hour) n=5 batches per group (two-way ANOVA Bonferroni post-test); **P<0.01 wild-type vs. VASP-KO. (**F**) Representative pictures showing the engulfed fluorogenic bioparticles in green and cell nuclei in red, bars=50µM.

Phosphorylation of the AMPK α-subunit is dynamically regulated by protein kinases and phosphatases. At least three kinases have been reported to phosphorylate AMPKα, namely, liver kinase B1 (LKB1), calcium-/calmodulin-dependent kinase kinase 2 (CaMKK2) and TGFβ-activated kinase 1 (TAK1). To elucidate the molecular mechanism underlying the impaired AMPK phosphorylation in VASP-deficient cells, kinase- and phosphorylation site-specific antibodies were used to investigated whether reduced expression and/or activation of the upstream AMPK kinases could account for the impaired AMPK phosphorylation in the absence of VASP. However, Western blotting of lysates from VASP-deficient macrophages did not reveal any obvious changes in the expression or activation of LKB1, CaMKK2, or TAK1 compared to wild-type cells (Figure 2). At least three protein phosphatases (PPs) have been reported to dephosphorylate and inactivate AMPK, namely PP1, PP2A and PP2C (Jeon, 2016). Mechanistically, PP1 is tightly associated with the protein phosphatase regulatory subunit 6 (PP1-R6), which physically interacts with the β-subunit of the AMPK complex and recruits PP1 to dephosphorylate the catalytic α-subunit at Thr-172 (Garcia-Haro et al., 2010). Interestingly, PP1-R6 contains a conserved peptide motif ^240^**FPVPP**FLL**EL**^249^ (numbering according to the human sequence, Figure 3A) that closely resembles the high-affinity EVH1 binding motif of ActA (Niebuhr et al., 1997).

**Figure 2.**
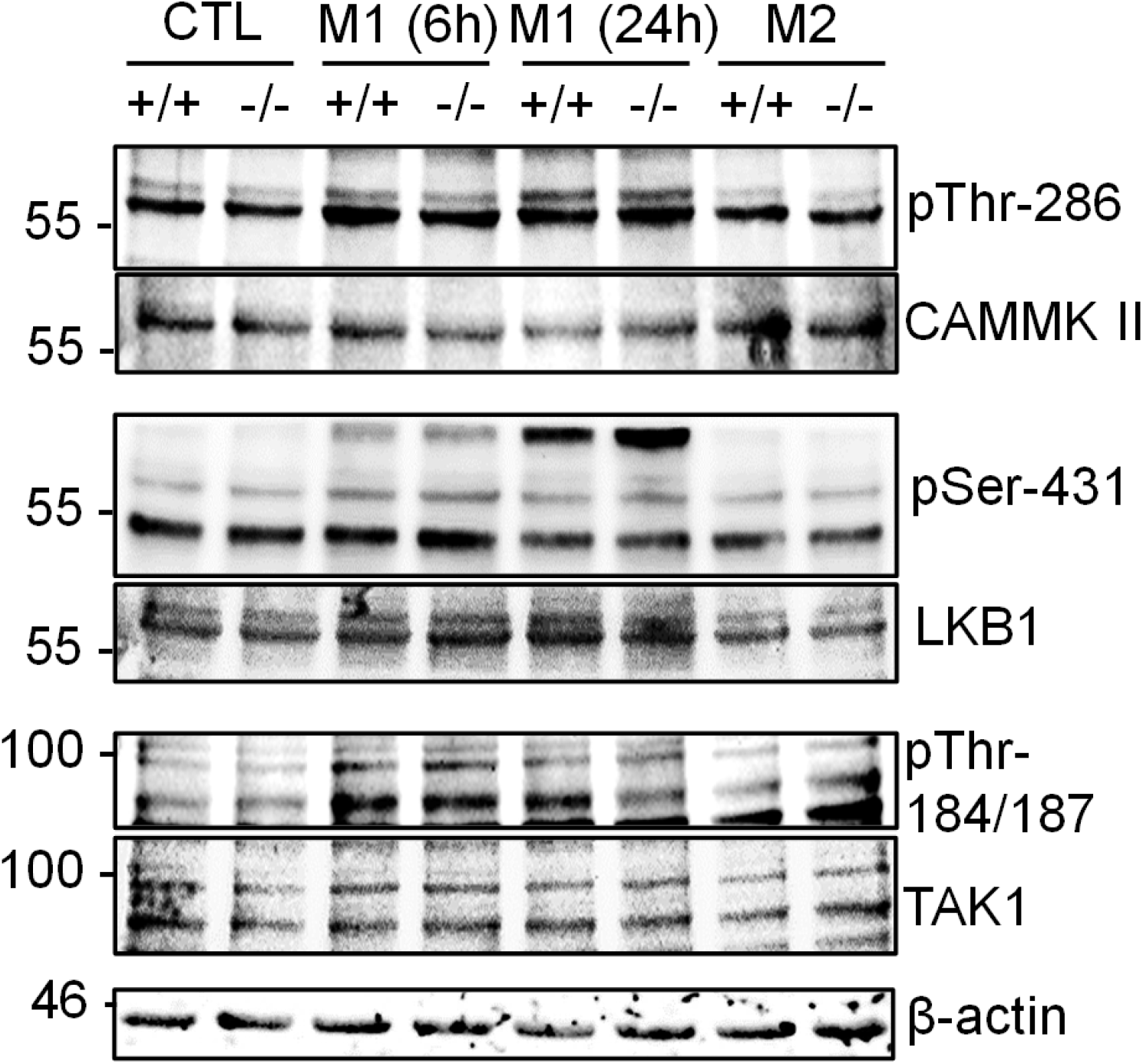
Representative Western blots showing the expression and phosphorylation (activation) of CAMMK II, LKB1 and TAK1 in wild-type (+/+) and VASP-deficient (-/-) macrophages.

**Figure 3.**
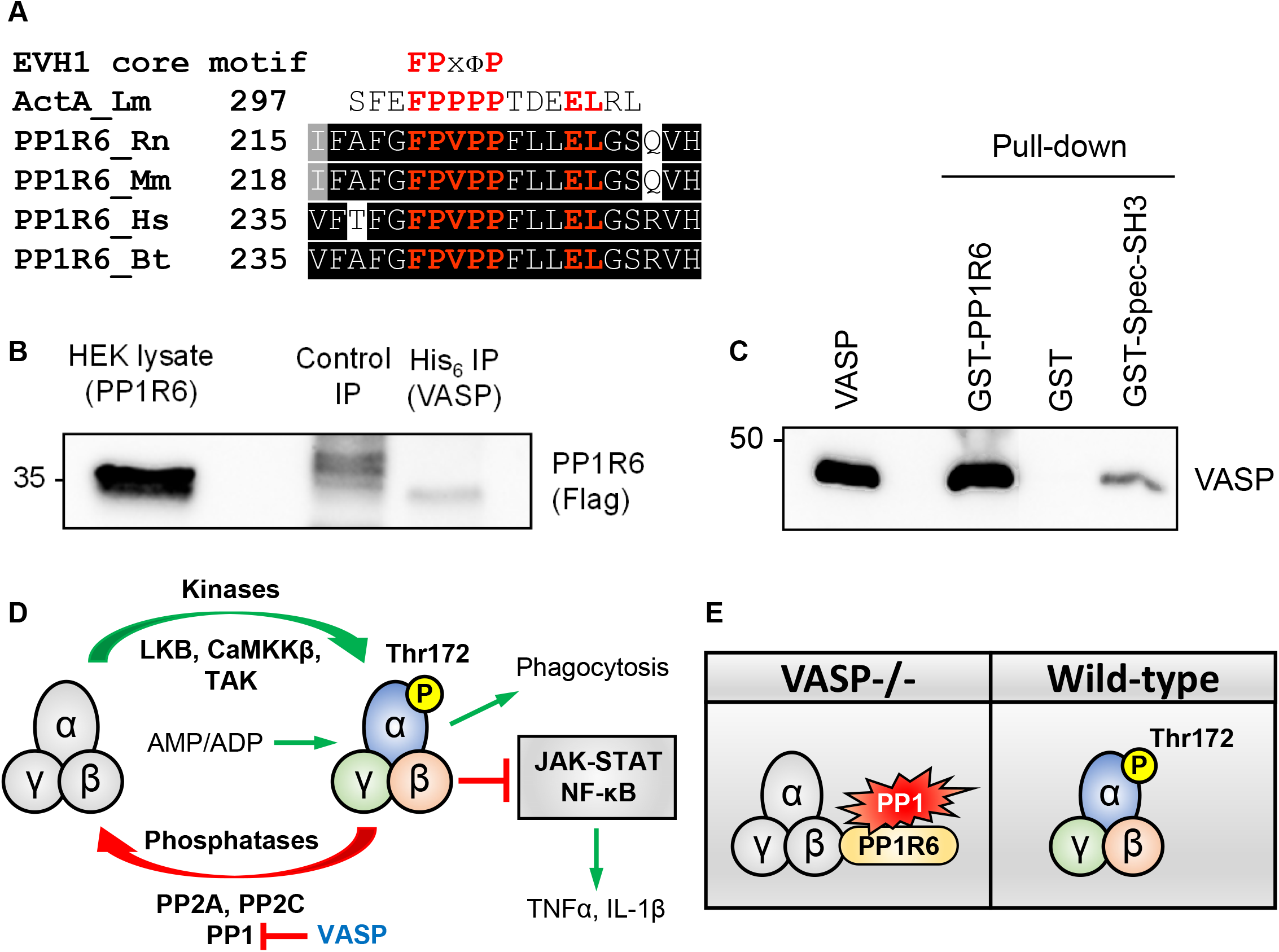
(**A**) Multi-sequence alignment of rat (Rn), mouse (Mm), human (Hs), and bull (Bt) PP1-R6 protein sequence. The putative VASP EVH1 domain binding motif is highlighted in red. The EVH1 core binding motif and the second, high-affinity EVH1 binding motif of listeria (Lm) ActA are shown for comparison (x, any amino acid; Φ, a hydrophobic amino acid). (**B**) Western blot indicating complex formation (immunoprecipitation) of VASP and PP1R6 in HEK cells transfected with His_6_-VASP and Flag-tagged PP1R6. (**C**) Recombinant VASP was incubated with equimolar amounts of the depicted, immobilized GST fusion proteins or GST alone (negative control). The precipitated material was analyzed with VASP-specific antibodies in Western blots, demonstrating direct binding of VASP to PP1-R6. Interaction with the immobilized GST-Spectrin-SH3 domain served as a positive control. (**D, E**) Regulation of AMPK activation by phosphorylation/ dephosphorylation on Thr172. Phosphorylation of AMPK at Thr-172 is induced by the upstream protein kinases LKB1, CaMKKβ and TAK1. Dephosphorylation of AMPK at Thr172 is regulated by protein phosphatases PP1, PP2A and PP2C. In macrophages, AMPK activation inhibits JAK-STAT and NF-κB signaling. This limits skewing macrophage function towards the M1 phenotype and hence reduces the expression of pro-inflammatory cytokines including TNFα and IL-1β. AMPK activity is also a driver for macrophage phagocytosis. AMPK phosphorylation is reduced in VASP-deficient macrophages. We hypothesize that VASP-binding to PP1-R6/PP1 limits the PP1-dependent de-phosphorylation of AMPK in wild-type cells. In the absence of VASP, AMPK dephosphorylation by the PP1-R6/PP1 complex is increased.

To determine whether a VASP and PP1R9 associated in living cells, HEK-293 cells were co-transfected with His_6_-VASP and a Flag-tagged PP1-R6 protein. Cells were lysed, immunoprecipitated with antibodies against VASP or control antibodies, and the precipitated material was probed with Flag-specific antibodies to detect PP1-R6. This revealed that VASP physically associated with PP1R6 (Figure 3B). To exclude that the binding of VASP to PP1R6 was indirect, the interaction of the purified proteins was analyzed in a cell-free system. In GST pull-down assays, immobilized GST-PP1-R6 readily bound to VASP, whereas equimolar amounts of GST alone showed no binding. Interestingly, GST-PP1R6 precipitated substantially more VASP than equal amounts of GST-Spectrin-SH3, which was used as positive control (Benz et al., 2008) (Figure 3C). Given that VASP itself is dephosphorylated by PP1 (Abel et al., 1995), it is tempting to speculate that loss of VASP increases PP1-R6 targeting to AMPK to promote kinase dephosphorylation, which would be expected to have implications on macrophage polarization and phagocytosis (Figure 3D and E). Since VASP is phosphorylated by AMPK (Blume et al., 2007), this may constitute a negative feedback loop that reduces AMPK activation in VASP-deficient macrophages, which in turn drives a STAT1/NF-κB mediated pro-inflammatory phenotype.

## Materials and Methods

Macrophage differentiation and polarization were previously reported (Laban et al., 2018). Briefly, BM-cells were isolated from wild-type and VASP^-/-^ mice, and collected by centrifugation at 300Xg for 5 minutes at 4°C. Erythrocytes were lysed with hypotonic buffer for 5 minutes, and washed with 1XPBS. BM-derived leukocytes were cultured in plastic culture dishes in basal RBMI-1640 containing 1X non-essential amino acid, 1X sodium pyruvate, MEM vitamins and P/S. After one hour, non-adherent cells were washed off and the medium was replaced with the basal RBMI-1640. Additionally, 10% FCS, 15 ng/ml M-CSF and 15 ng/ml GM-CSF were added to the culture medium. After 7 days, the macrophages were ready for polarization. To induce pro-inflammatory M1 macrophages, BM-derived macrophages were treated with 1 ng/ml IFNγ and 10 ng/ml LPS. Meanwhile, alternative M2 macrophages were polarized after treatment with 25 ng/ml IL-4. mRNA levels of TNFα and IL-1β were determined as previously described (Laban et al., 2018).

Phagocytosis assay. BM-derived macrophages (2×10^5^ cells/ 100 µL 1X Opti-MEM medium) were plated in a 96-well plate (Essen Biosceince ImageLock, Germany). Afterwards, the macrophages were then incubated with homogeneous pHrodo™ Green *E. coli* BioParticles® (#P35366, Thermofisher, Germany) at a final concentration 1 mg/ml for further 5 hours using live cell imaging (IncuCyte, Essen Bioscience) to measure the relative fluorescence intensity. The % Net Phagocytosis was calculated by subtracting the no-cell control from the average fluorescence intensity.

For immunoprecipitations with tagged proteins, HEK cells were co-transfected with human flag-tagged PP1R6 (Origene) and His_6_-tagged VASP (Benz et al., 2009) using Lipofectamine 2000 (Invitrogen, Karlsruhe, Germany). Cells were lysed in buffer A (40 mM HEPES-NaOH, pH 7.5, 100 mM NaCl, 1 % Igepal CA-630, and protease/phosphatase inhibitor cocktails). Lysates were clarified by centrifugation for 10 minutes at 20,000 Xg at 4 °C and VASP was subjected to immunoprecipitation using either His_6_- specific epitope tag affinity matrix (Biolegend #900501) or control antibody matrix (Novagen #69026). Precipitated material was probed by Western blotting with anti-Flag antibodies (Sigma-Aldrich).

GST pull-down assays were performed as described (Benz et al., 2016). Briefly, 500 ng of purified His_6_-VASP (Benz et al., 2009) was incubated with 5 μg GST-PP1R6, equimolar amounts of GST alone (negative control), or GST-Spectrin-SH3 (positive control (Benz et al., 2008)) coupled to glutathione- Sepharose (GE-Healthcare, Freiburg, Germany) in buffer A supplemented with 25 ng/μl BSA to block non-specific interactions. After extensive washing, the precipitated material was analyzed by Western blotting with anti-VASP antibodies (Benz et al., 2008).

## Acknowledgements

This work was supported by the Deutsche Forschungsgemeinschaft (SFB 834/A8 to PMB, SFB 834/A5 to IF). PMB was also supported by the German Center for Cardiovascular Research (DZHK B14-028 SE). The authors are indebted to Andreas Weigert for scientific input and assistance.

